# Connectomic analysis reveals the axial circuit for self-righting posture control in *Drosophila*

**DOI:** 10.64898/2026.07.25.740718

**Authors:** J. Picao-Osorio, W. Roseby, C.E. Hancock, A. Cardona, C.R. Alonso

## Abstract

Postural control, the capacity of an animal to detect and correct an inappropriate body orientation, is a widespread and fundamental neurobiological function, yet the neural circuits that implement it remain poorly resolved in most species. The larva of *Drosophila melanogaster* performs a posture control stereotyped self-righting (SR) manoeuvre when turned upside-down, a behaviour previously shown to depend on a pair of segmentally repeated lateral transverse motor neurons (LT1/2-MNs) and on the normal expression of several Hox-targeting microRNAs. Here, we use a connectomics and a functional behavioural approach to trace and test the sensory, interneuronal and motor architecture of the SR circuit along the antero-posterior axis. Starting from the LT1/2-MNs, we identify a set of pre-motor interneurons, their principal upstream partners, and a population of class IV multidendritic sensory neurons as core components of the circuit, and show, through neuron-specific thermogenetic silencing, that inhibiting the great majority of these elements significantly impairs SR performance. We further find substantial overlap between the SR circuit and the previously described nociceptive rolling and touch crawling circuits, converging on shared interneurons including DnB, A02o (Wave-1) and TePn05. Network and axial connectivity analyses reveal a nested, hub-like organisation, a small number of integrator neurons bridging sensory and motor sub-networks, and a consistent decline in synapse number and density towards posterior segments. Together, these data yield the first axial wiring diagram for a postural control circuit in any animal and offer a set of structural principles including: hub organisation, shared sensory-motor structure, and antero-posterior connectivity gradients, which may extend to other segmentally organised nervous systems.

## INTRODUCTION

Most animal forms display the capacity to control their body posture and ensure a suitable orientation, be it regarding to the polarity of the gravitational field (Benzer, 1967) or in respect to the substrates they inhabit (Ashe, 1970; Faisal and Matheson, 2001). This orientational preference possibly stems from anatomical asymmetries – for instance, along the dorso-ventral axis – selected as adaptations to biological functions. For instance, ventral walking appendages or dorsal camouflage are of negligible value if presented in the wrong orientation. Accordingly, when the animal finds itself in an unsuitable position, it quickly does something about it by promoting what comes across as a rather complex set of estimates, computations and actions aimed at restoring the right posture. This succession of activities is what we shall term postural control.

The mere existence of a postural control system implies an internal ability to: (i) monitor the relative (or absolute) position of the body at a given time, (ii) compare values against an internally pre-conceived setting, and (iii) in the event that discrepancies exceed an acceptable tolerance, (iv) trigger a motor programme able to modify body posture, and (v) eventually halt postural adjustment once a suitable orientation is restored. Altogether, this comes across as a formidable illustration of a neurobiological control mechanism functionally coupled to the existence of morphological asymmetries showcasing the high level of interdependence between form and function in animal systems. Despite their apparent complexity, postural control systems can be found in complex animals (fish, amphibians, reptiles and mammals)(Arabyan and Tsai, 1998; Ashe, 1970; Bisazza et al., 1996; Pellis et al., 1991; Webb, 2002), as well as in simpler animal forms (nematodes and insects)(Crisp et al., 2008; Hums et al., 2016). For example, the ability of human babies to roll and rectify their position should they be put ‘on their backs’ is a common medical milestone used to assess motor development (McGraw, 1941). Similarly, the larval form of the fruit fly *Drosophila melanogaster* is capable of postural control via an innate motor sequence termed self-righting (SR) which allows the larva to correct its position if turned upside-down (Figure 1A) (Issa et al., 2019; Picao-Osorio et al., 2015). Interestingly, previous genetic experiments conducted in our laboratory revealed that normal expression of dozens of non-coding RNAs (microRNAs, miRNAs) is necessary for the fly larva to perform a normal SR response (Klann et al., 2021; Picao-Osorio et al., 2017) suggesting a genetic basis to SR. In addition, molecular pathway analyses revealed that many of the SR-related microRNAs target the Hox genes (Picao-Osorio et al., 2017, 2015), which encode a family of developmental regulators with specialised roles along the head-to-tail (antero-posterior) axis (Mallo and Alonso, 2013). Interestingly, behavioural observations strongly suggest a differential recruitment and sensory properties of larval segments during the SR sequence (Loveless et al., 2021; Picao-Osorio et al., 2015; Roseby et al. 2026) suggesting that the underlying sensory motor processes involved in SR may themselves vary along the main body axis. To advance the understanding of the mechanisms by which the genetic system influences SR behaviour, it is therefore necessary to map the neural circuitry that enables the SR response not only in one given segment but along the main axis of the animal.

**Figure 1.**
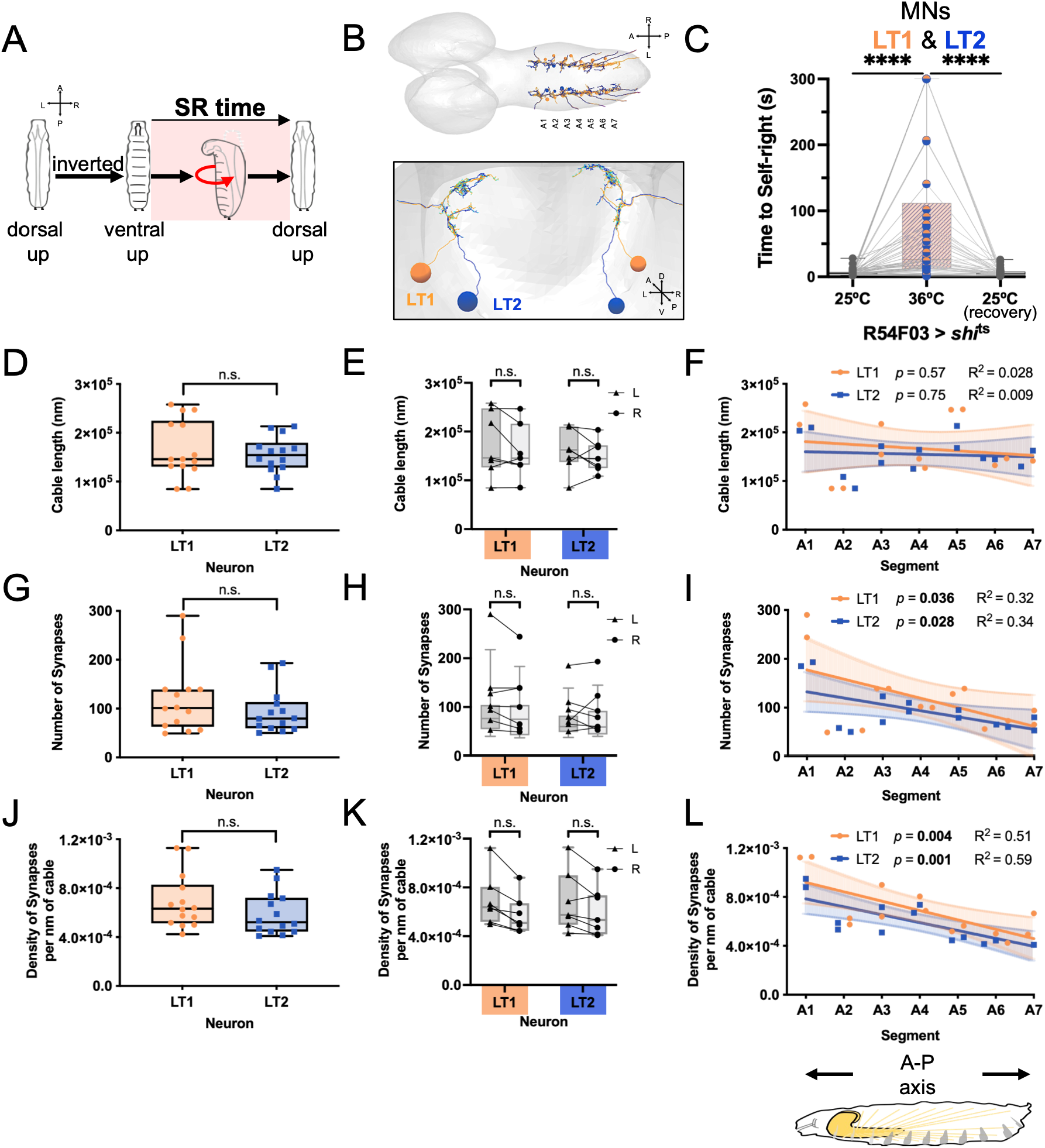
Neuronal morphology of the motor neurons LT-1/2 along the anteroposterior axis. **(A)** Conceptual diagram of the Drosophila larval self-righting behaviour. **(B)** Upper panel, first instar larval central nervous system with the neuronal reconstructions of LT-1 (orange) and LT-2 (dark blue) MNs along abdominal segment A1 to A7. Anterior is to the left. Lower panel, transverse view of the motor neuronal reconstructions of LT-1/2 in segment A2. **(C)** Thermogenetic inhibition of the neural activity in the motor neurons LT-1 and LT-2 (R54F03>*shi* ^ts^) (n = 54 larvae; \*\*\*\**p*<0.0001, Wilcoxon matched-pairs test) and SR assay. Boxplots with minimum and maximum values, and individual data points of larvae tested initially at the permissive temperature of 25°C, at the restrictive temperature of 36°C, and the recovery permissive temperature of 25°C. **(D)** Quantification of the nanometer (nm) cable length in LT-1 and LT-2 in abdominal segments A1 to A7 (n=14 LT1 and 14 LT2 MNs; n.s. *p*>0.05, Welch’s *t*-test). Boxplots with individual data points. **(E)** Quantification of the cable length in the Left and Right sides of LT-1 and LT-2 in abdominal segments A1 to A7 (n=7 for each side and MN; n.s. *p*>0.05, paired *t*-test). **(F)** Quantification of cable length in LT-1 and LT-2 along the abdominal segments A1 to A7 [slope n.s. *p*>0.05 and goodness of fit (R^2^), simple linear regression]. **(G)** Quantification of the number of upstream synapses in LT-1 and LT-2 in abdominal segments A1 to A7 (n=14 LT1 and 14 LT2 MNs; n.s. *p*>0.05, Mann-Whitney *U* test). Boxplots with individual data points. **(H)** Quantification of the number of upstream synapses in the Left and Right sides of LT-1 and LT-2 in abdominal segments A1 to A7 (n=7 for each side and MN; n.s. *p*>0.05, Wilcoxon matched-pairs test). **(I)** Quantification of number of upstream synapses in LT-1 and LT-2 along the abdominal segments A1 to A7 [slope * *p*<0.05 and goodness of fit (R^2^), simple linear regression]. **(J)** Quantification of the synaptic density (number of upstream synapses per nanometer of cable length) in LT-1 and LT-2 in abdominal segments A1 to A7 (n=14 LT1 and 14 LT2 MNs; n.s. *p*>0.05, Mann-Whitney *U* test). Boxplots with individual data points. **(K)** Quantification of the synaptic density in the Left and Right sides of LT-1 and LT-2 in abdominal segments A1 to A7 (n=7 for each side and MN; n.s. *p*>0.05, paired *t*-test). **(L)** Quantification of the synaptic density in LT-1 and LT-2 along the abdominal segments A1 to A7 [slope *** *p*≤0.001 and goodness of fit (R^2^), simple linear regression].

In this study, we apply a neural connectomics approach to determine the identity of the neural substrates underlying SR behaviour in the *Drosophila* larva. Using as a starting point a pair of metameric motor neurons (i.e. lateral transverse motor neurons LT-MNs 1/2) identified in previous work (Picao-Osorio et al., 2015; Zwart et al., 2016), we trace back the interconnected pre-motor and interneuronal networks and proceed to map the sensory elements that mediate this behaviour. Perhaps uniquely, our work exploits the natural repeating structural nature of the larval nervous system to analyse the segmentally dependent cellular modifications of the circuitry along the main body axis. This comprehensive analysis led to the generation of a wiring diagram for the sensory, interneuronal and motor command elements underlying SR in the fruit fly larva. In addition, functional experiments involving thermogenetic inhibition of the individual components of the SR network provide the basis to convert the SR wiring diagram into the first version of the neural circuit that enables SR. Our comparison of the SR neural architectures along the antero-posterior axis determines the patterns of connectivity within and across individual segments observing how the stereotyped array of SR neurons is modified according to axial position allowing us to deduce a set of principles of how the network might mediate the postural detection and permit the execution of this behaviour.

The work generates the first draft of an axial neural circuit involved in postural control in a metazoan and suggests a set of rules for information flow and antero-posterior architecture that might conform a point of reference for the analysis of similar circuits in other animal systems. We speculate that our work might also aid the design of future control systems for soft body robotic units.

## MATERIALS AND METHODS

### Drosophila maintenance and stocks

Flies were reared following standard procedures at 25°C on a 12-hr light/dark cycle and 50– 60% relative humidity. We used the following *Drosophila melanogaster* strains that were obtained from Bloomington *Drosophila* Stock Center (BDSC) or were kind gifts from the laboratories of Marta Zlatic (MRC LMB, Cambridge, UK), Matthias Landgraf (University of Cambridge, UK) and James W. Truman (University of Washington, Seattle, USA): *w^1118^* (BDSC #5905); R54F03-GAL4 [MN-LT-1/2 (Picao-Osorio et al., 2015), BDSC #39078, (Manning et al., 2012)]; pre-LT GAL4 drivers from (Zwart et al., 2016) A18j (SS01970-GAL4, BDSC #603375), A01c (R75H04-GAL4, BDSC #39909), A14a (SS01411-GAL4, BDSC #604309), A19l (R16E12-GAL4, BDSC #48730) and A26f [SS04290-GAL4 (Meissner et al., 2025), BDSC #603814]; A09l [DnB (Burgos et al., 2018), 412-GAL4 (PB[IT.Gal4]0412) (Gohl et al., 2011)], A19f (SS24070-GAL4, BDSC #603098); rolling (Ohyama et al., 2015) md class IV (*ppk*1.9-GAL4 (Ainsley et al., 2003), chordotonal organs (*iav*-GAL4, BDSC #52273), Basins (R72F11-GAL4, BDSC #39786), A05q (R47D07-GAL4, BDSC #50304), A23g (SS02139-GAL4 (Jovanic et al., 2019), BDSC #60256), Goro (R16E11-GAL4, BDSC #48729), TePn05 (R61A01-GAL4, BDSC #39269); UAS-*shi*^ts^ [(Kitamoto, 2001) BDSC #44222)

#### Behavioural experiments

Thermogenetic experiments to inhibit SR neurons during larval self-righting behaviour were conducted similarly as previously described (Picao-Osorio et al., 2015). Briefly, we crossed parental flies of the neuronal-specific GAL4 drivers and the UAS-*shi*^TS^ to inhibit neural communication at the restrictive temperature of 36°C. These parental crosses were kept on small collection cages with apple juice agar plates supplemented with yeast paste. Embryos were collected from these plates and aged until stage 17. Freshly hatched first instar larvae (<30 minutes post-hatching) were placed on a 1.5% agar substrate (2mm thickness) onto a custom-built temperature controller-Peltier module adjusted to the permissive temperature of 25°C and were allowed to acclimatise for 1 minute. We tested the same larva at three temperatures. First, at the permissive temperature of 25°C, larvae were gently rolled over using a single-bristled paintbrush to an inverted position (ventral denticle belts up) and the time taken by the larvae to self-right –dorsal longitudinal trachea up– was measured. Second, the temperature was raised to the restrictive 36°C to inhibit neural activity for 2 minutes and tested for SR behaviour. Third, temperature was lowered back to the 25 C° and larvae were allowed to recover for 5 minutes before being rolled onto their dorsal sides and allowed to self-right for a third time (recovery test). SR times were measured to a maximum of 5 minutes, after which the assay was stopped and larvae rolled back to their original position. A minimum of 20 larvae and a maximum of 54 larvae were tested per genotype (see figure legends for details). All experiments were done in a room maintained at 25 C°. Plots and statistical tests were performed in GraphPad (v. 11.0.2) using a Wilcoxon matched-pairs signed-rank test.

#### Circuit reconstruction

We reconstructed neuronal morphologies and synapses on a serial section transmission electron microscopy volume of a complete first-instar larval central nervous system (Ohyama et al., 2015). Neuron reconstructions were performed using CATMAID [Collaborative Annotation Toolkit for Massive Amounts of Image Data, (Saalfeld et al., 2009)] using iterative reconstruction methods (Schneider-Mizell et al., 2016). Annotation of synapses followed the previously described criteria of mature synapses, including a dark, T-shaped synaptic cleft, pre/post-synaptic vesicles and the common presence of a mitochondrion near the pre-synaptic site (Eschbach et al., 2021; Ohyama et al., 2015; Schneider-Mizell et al., 2016). Motor neurons LT-1 and LT-2 and preLT INs were identified and reconstructed in abdominal segments based on the position of their cell bodies, axonal and dendrite projection patterns (Landgraf et al., 1997; Zwart et al., 2016). Furthermore, we used (and extended in some segments) the previously reconstructed neuronal elements of the rolling and peristaltic locomotion circuits (Ohyama et al., 2015; Takagi et al., 2017). Our reconstructed SR circuit is composed of the following axial neurons: *LT-1* and *LT-2* in abdominal segments A1 to A7; *A26f* in A1 to A6 (right side of A5 was not reconstructed); *A18j* in A1 to A6 (left side of A6 was not reconstructed); *A01* and *A14a* in A1 to A6; A19l in A1 to A3 (left side of A3 was not reconstructed); *A09l* in A1 to A5; *A27k* in A1, A2, A3 left, A4 right, A5 right; *A19f* in A1 to A6; *md calss IV* in A1 to A6 (ddaC in A1-A5, A6 left; v’ada in A1-A4, A5 right; vdaB in A1-A6, except A5 right); *chordotonal organs* in A1-A6 (lch5-1 in A1, A2, A4, A5 left; lch5-3 in A1, A2, A4 left, A5 right; lch5-5 in A1, A2, A4 right, A5 right, A6 right; lch5-2/4 in A1, A2 left, A3, A4, A5 left, A6); *A09a (Basin-2)* in A1-A4 (except A3 left); *A09g (Basin-3)* and *A09c (Basin-4)* in A1-A4; *A05q* and *A23g* in A1-A2; *Goro* in T3; *TePn05* in A8; and *A02o (Wave-1)* in A1 to A5 and A6 left.

#### Circuit Analysis

After annotating and reconstructing the abdominal neurons of the SR circuit, we exported for each neuron along the A-P axis (on both sides) neuronal features such as raw and smooth cable length, number of input/output synapses for all pre-and postsynaptic partners. Since reconstructions had been annotated over time by different contributors, we compiled the neuronal CATMAID outputs of every source neuron by their identity/name, body segment and side, using a custom Python pipeline (pandas and openpyxl libraries). Then, compiled upstream and downstream data was filtered for the SR neurons and summarised for each neuron along the body segments and side, also using a custom Python pipeline. The neurons belonging to the two sensory groups were pooled together (i.e. ddaC, v’ada and vdaB as mdIV; and lch5-1/3/5/2–4 as “chordotonals”) for the overall connectivity analysis (not in the axial and lateral polarity analysis, see below). Within each group, the median proportion of a source neuron’s total synapses was taken across all individual instances — segmental homologous and left/right sides. Plots and statistical tests of the quantification of neuronal cable length, number, density and proportion of synapses were accomplished in GraphPad (v. 11.0.2) using Welch’s *t*-test and paired *t*-test for normally distributed data, and Mann-Whitney *U* test, Wilcoxon matched-pairs test and Kruskal-Wallis test with Dunn’s multiple comparisons test for not Gaussian data. Normality was accessed with Shapiro-Wilk test. Furthermore, to test whether the neuronal features changed along abdominal segments, we performed simple linear regressions and assessed if the slope of the regression was statistically significant from zero.

Neuronal synaptic connectivity, hierarchical clustering and network analysis were performed using R (v. 4.6.0)(Team, 2026) in Rstudio (v. 2026.05.0) (team, 2026) using the tidyverse and readxl packages for data structuring and ggplot2 for data visualisation. Neuronal connectivity was depicted at the level of individual source neurons as fan diagrams: each focal neuron was placed at the centre of a semicircular arrangement of its downstream partners, with arrow width encoding connection strength (median proportion, binned into discrete categories) and the median and range across segmental/left-right instances annotated alongside each arrow (using the purrr, ggplot2 and scales packages). For the hierarchical clustering and network analyses, the group-level medians were arcsine square-root transformed to stabilise the variance across the wide range of observed proportions. Hierarchical clustering of the transformed group-level connectivity matrices was shown as a clustered heatmap using Euclidean distance and average linkage (pheatmap package). Dendrogram branches were rotated at existing merge points, without altering cluster topology, to approximate the expected sensory-to-motor circuit hierarchy. A force-directed network graph was generated (igraph, ggraph, and tidygraph packages) to provide a conceptual overview of circuit connectivity. Source and target groups connected were represented as nodes, joined by directed, curved edges indicating the direction of connectivity. Node position was determined using the Fruchterman-Reingold algorithm, with vertical position additionally constrained so that primary sensory neurons (class IV multidendritic neurons, mdIV; chordotonal neurons) were positioned at the top of the network and motoneurons (LT1, LT2) at the bottom, reflecting the expected sensory-to-motor direction of information flow; horizontal position within each hierarchical tier remained determined by the force-directed layout.

Connectivity polarity was analysed for each synaptic connectivity wiring diagram edge in two measures of connection relative to the source neuron: (i) axial polarity (Anterior / Same segment / Posterior), from the partner’s segment, and lateral polarity (Ipsilateral / Contralateral), from its side. Synapse-weighted percentages of each category, summarised by neuron group and connection direction (upstream/downstream), are shown as mirrored bar charts overall.

Code for data compilation, filtering, summarisation and connectivity was developed with the assistance of the AI model Claude (Claude Sonnet 4.5; Anthropic, San Francisco, CA, USA), under the authors’ direction and with manual verification of all outputs. All code developed and implemented is available via email request to the corresponding authors.

## RESULTS AND DISCUSSION

### Neuroanatomical analysis of MN-LT1/2 system along the antero-posterior axis

Previous work, conducted with the goal of mapping the cellular site of action of genes necessary for SR (Picao-Osorio et al., 2015), identified that a pair of metameric motor neurons (MN), called the LT1/2-MNs (Landgraf et al., 1997) as the first cellular component required for normal SR, whereby their neural activity inhibition leads to a significant increase in SR time [(Picao-Osorio et al., 2015) and **Figure 1C**]. Through the combination of gene expression and functional imaging analyses this work highlighted that individual abdominal segments in the larval ventral nerve cord (VNC) might play distinct roles in regard to the SR sequence (Picao-Osorio et al., 2015) ascribing an important role to a series of abdominal segments (A2-A5). To advance the understanding of the neural substrates underlying SR we applied a neural connectomics approach by which the connectivity patterns of individual larval neurons can be reconstructed from inspection of a serial electron microscopy ultrastructural volume of a reference first-instar larval nervous system (Ohyama et al., 2015; Schneider-Mizell et al., 2016). A previous study had already inspected the upstream connectivity patterns of the LT1/2 MNs in the first abdominal segment A1 (Zwart et al., 2016) and identified a group of excitatory and inhibitory interneurons providing essential information to the LT1/2 MNs for normal larval locomotion. Building on these observations, we firstly reconstructed the LT1/2 MNs across the entire abdominal region, documenting the properties of the LT1/2 MN system in each abdominal segment (Figure 1). A first question that emerged during this characterisation was whether the individual LT1-and LT2-MNs had different intrinsic properties that might be relevant for their roles in SR control. To explore this question, we used CATMAID to assess the level of anatomical similarity across MN-LT1 and MN-LT2 types by quantitatively analysing two key metrics: (i) dendritic arbour cable length; and (ii) upstream synapses, previously determined to be the two most effective measures for anatomical comparisons across neural types (Schneider-Mizell et al., 2016). Quantification of cable length for each neuron across the larval abdomen show that both motor neurons MN-LT1 and MN-LT2 possess similar values of cable length (**Figure 1D**) and there is no significant difference between the left and right sides (**Figure 1E**). Furthermore, mapping cable length to individual abdominal segments does not show inter-neuronal differences between MN-LT1 and MN-LT2 (**Figure 1D**) or apparent differences along abdominal segments, with segments A1 and A5 displaying the highest cable length values for both motor neurons (**Figure 1F**). Analysis of input synapse numbers showed no clear differences in either between MN-LT1 and MN-LT2 (**Figure 1G**) or the left and right sides of each MN (**Figure 1H**). Interestingly, we observed a significant decrease in the number of synapses for both MN-LT1/2 towards the posterior abdominal segments (**Figure 1I**), while both MNs display very similar properties along the antero-posterior axis, with A1 showing ≥2-fold number of synapses than those present in the other abdominal segments for both neurons. Next, we analysed the synaptic density by calculating the ratio of synapse number per nanometre of cable length and found no difference between the MNs (**Figure 1J**) or between the left and right sides of each MN (**Figure 1K**) Remarkably, we observed an axial trend with a decreasing synaptic density for MN-LT1 and MN-LT2 as you move from anterior to posterior abdominal locations (**Figure 1L**). All in all, these data argue that (i) that the MN-LT1/2 motor neurons have very similar anatomical properties along the AP axis suggesting that they may possess very similar or equivalent functions; and (ii) that their density of synapses of these two neurons decreases towards the posterior regions of the VNC. The observed decrease in synapses of MN-LT1/2s might mean that the function of these neurons is less significant towards the posterior end of the larval body; these data also suggest that to achieve a comparable level of activity these neurons will require a higher level of stimulation from the pre-motor system or possess a lower threshold of activation to exert their effects on the muscle system.

### Pre-motor wiring of MN-LT1/2s along the main body axis

As mentioned above, previous work had identified the basic pattern of innervation to the MN-LT1/2 motor neurons for the first abdominal segment (A1) (Zwart et al., 2016). In order to develop an axial map of pre-motor connectivity to the MN-LT1/2 system we used the available A1 information to produce a subset of candidate pool of pre-motor neurons, including the excitatory (eIN-1, eIN-2 and eIN-3) and inhibitory (iIN-1, iIN-2 and iIN-3). Of these we noted that only eIN-1 (A18j), eIN-2 (A01c), iIN-1 (A14a) and iIN-3 (A19l) passed the standard ≥3 synapse threshold of connectivity associated with bona fide functional neuronal links (Schneider-Mizell et al., 2016). This analysis also revealed that interneuron A26f was also recruited as a pre-motor element to the MN-LT1/2 system (**Figure 2A**). Notably, a salient aspect of pre-motor organisation that emerged from this analysis is that all pre-motor components appear contralateral to MN-LT1/2 indicating that information for motor neuron modulation is collected from the opposite side of the body to the motor neuron site (**Figure 2A**).

**Figure 2.**
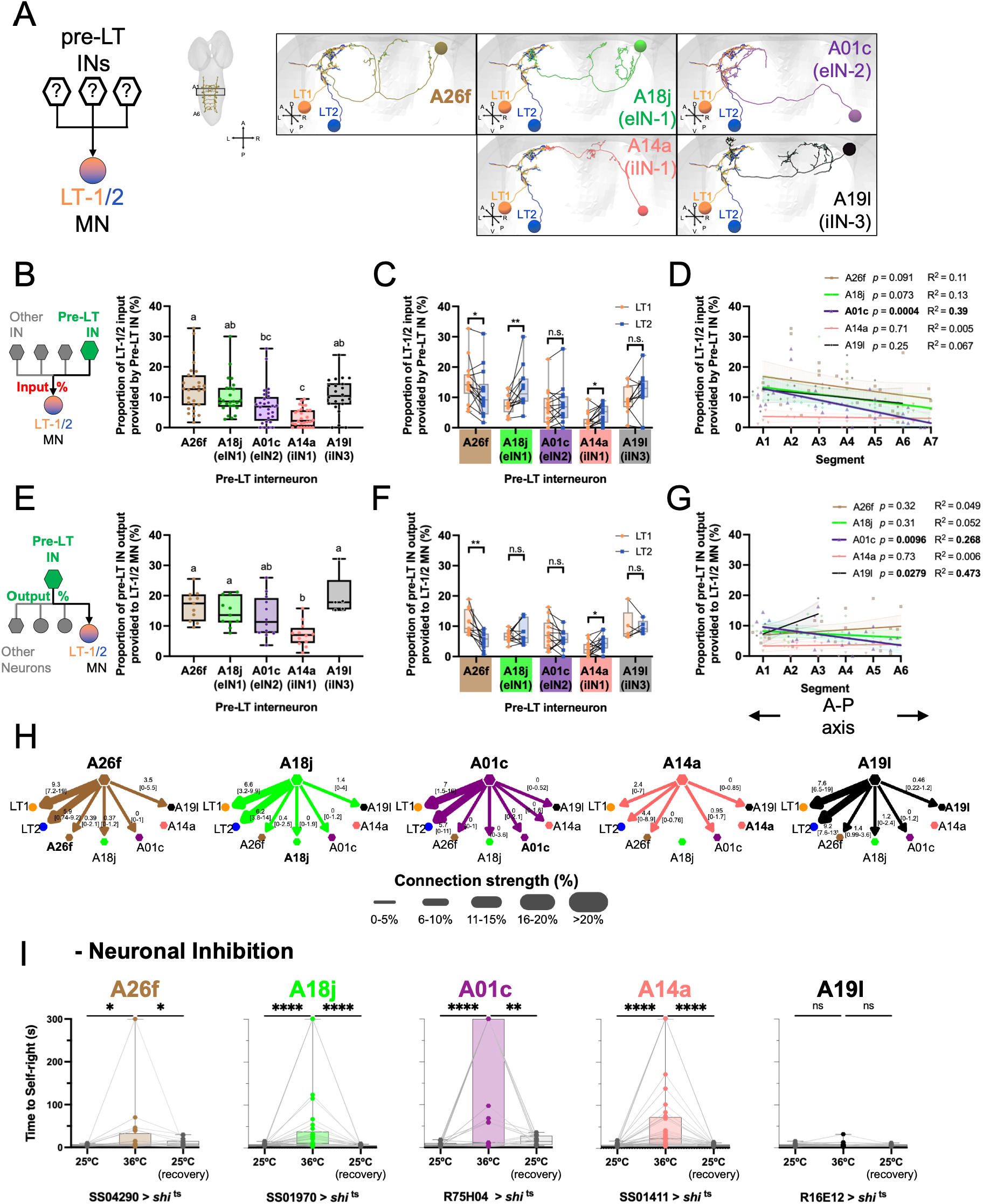
Premotor interneurons of LT-1/2 along the A-P axis. **(A)** Abdominal neuronal reconstructions of premotor interneurons (IN) of LT1/2: A26f (brown), A18j (green), A01c (purple), A14a (salmon) and A19l (black). Right panels, transverse views of the pre-LTs INs and the MNs LT-1 (orange) and LT-2 (dark blue) in segment A2. **(B-D)** Percentage of LT-1/2 upstream connectivity provided by the pre-LTs INs in abdominal segments A1 to A7. (B) Boxplots with individual data points of the 28 LT-1/2 MNs, bilateral MNs in 7 abdominal segments. Boxplots with different letters indicate significant differences (Kruskal-Wallis test with Dunn’s multiple comparisons test, n.s. *p*>0.05, \*\**p*<0.01). (C) Quantification of the Percentage of LT-1 and LT-2 upstream connectivity provided by the pre-LTs INs (n=14 for each LT-MNs; n.s. *p*>0.05, \**p*<0.05, \*\**p*<0.01, Wilcoxon matched-pairs test), and (D) along the abdominal segments A1 to A7 [slope n.s. *p*>0.05, \*\*\**p*<0.001 and goodness of fit (R^2^), simple linear regression]. **(E-G)** Percentage of pre-LT IN downstream connectivity provided to the LT-MNs in abdominal segments A1 to A6. (E) Boxplots with individual neuronal data points of 11 A26f (from A1 to A6, right side of A5 was not reconstructed), 11 A18j (from A1 to A6, left side of A6 was not reconstructed), 12 A01 and A14a (from A1 to A6), and 5 A19l (from A1 to A3, left side of A3 was not reconstructed). Boxplots with different letters indicate significant differences (Kruskal-Wallis test with Dunn’s multiple comparisons test, n.s. *p*>0.05, \**p*<0.05). (C) Percentage of pre-LT IN downstream connectivity provided to LT-1 and LT-2 (n.s. *p*>0.05, \**p*<0.05, \*\**p*<0.01, Wilcoxon matched-pairs test), and (D) along the abdominal segments A1 to A6 [slope n.s. *p*>0.05, \**p*<0.05, \*\*\**p*<0.001, and goodness of fit (R^2^), simple linear regression]. (**G**) Fan diagrams displaying the downstream connectivity of each pre-LT IN to the LT-1/2 MNs and the pre-LT INs. The thickness of the arrows represents the median downstream percentage connectivity with respective values, and minimum and maximum values in squared brackets. (**H**) SR assay and thermogenetic inhibition of the neural activity in the pre-LT INs A26f (SS04290>*shi* ^ts^; n=20 larvae), A18j (SS01970>*shi* ^ts^; n=33 larvae), A01c (R75H04>*shi* ^ts^; n=44 larvae), A14a (SS01411>*shi* ^ts^; n=36 larvae), A19l (R16E12>*shi* ^ts^; n=34 larvae). Boxplots with minimum and maximum values, and individual data points of larvae tested initially at the permissive temperature of 25°C, at the restrictive temperature of 36°C, and the recovery permissive temperature of 25°C (n.s *p*>0.05, \**p*<0.05, \*\**p*<0.001, \*\*\*\**p*<0.0001, Wilcoxon matched-pairs test). (**I**) Connectivity network of the MNs LT-1/2 and upstream INs. The thickness of the arrows depicts the median downstream percentage connectivity.

A survey of the percentage of LT-1/2 upstream connectivity provided by the pre-LTs INs in abdominal segments A1 to A7 shows that the MN-LT1/2s have the highest level of input connectivity from the A26f interneuron (median 13% of inputs) (**Figure 2B**) with a significantly higher input in LT-1 (**Figure 2C**). In contrast, the MN-LT1/2s system receives the least input from A14a (iIN-1; median of 3%) with the other three interneural components showing intermediate levels of connectivity (median 7-10%) (**Figure 2B**), and A18j (eIN1) having a significantly higher input in LT-2 (**Figure 2C**). Axial mapping of these patterns of connectivity reveals that A01c (eIN2) present a significant decrease in its level of input that runs from anterior to posterior sites (**Figure 2D**). The pre-LTs A26f, A18j and A19l display similar, but not significant, negative trends towards posterior, whereas A14a (iIN-1) shows a uniform trend along the AP axis (**Figure 2D**).

Output proportionality connectivity analysis of the pre-motor elements shows that A26f, A18j, and A01c INs all display ≥13% median of their outputs linked to the MN-LT1/2s system (**Figure 2E and 2H**); A14a shows a much-reduced level of output towards MN-LT1/2s (median ∼7%), whilst A19l displays an apparently higher level of output links committed to the MN-LT1/2 system (∼20%). However, the significance of this latter observation on A19l is unclear given that our axial analysis could only be reliably identify this neuron across segments A1-A3 (i.e. seen on left and right sides in A1-A2, and only on the right-hand side in A3). Also, the pre-LT INs show a moderate (∼3%) to weak connectivity among themselves (**Figure 2H**). Furthermore, axial analysis reveals a significant decrease in A01c output to LT-1/2 MNs towards the posterior abdominal segments (**Figure 2G**).

To complement our connectomics and neuronal reconstruction analyses, we carried out a series of functional tests aimed at establishing the roles of the pre-motor components described above in SR behaviour. For this we systematically suppressed the roles of each pre-motor element expressing a dominant negative temperature-sensitive form of *shibire* (*shi*) (Kitamoto, 2001) to inhibit synaptic vesicle recycling in a neuron-specific manner via the UAS-Gal4 binary system (Brand and Perrimon, 1993). Neuron specific drivers for A18j, A01c, A14a, and A19l were previously described (Zwart et al., 2016); for manipulations of the A26f interneuron a newly published split-Gal4 driver was employed [SS04290, J. W. Truman personal communication; (Meissner et al., 2025)] which results from the overlapping activities of VT045148-AD ∩ R22E06-DBD. The results of these experiments indicate that when all these pre-motor interneuronal elements are inhibited this leads to a significant impact on SR behaviour manifested by an increased SR time, an indication of poor postural control performance, with the exception of interneuron A19l (iIN-3) which shows no effect (**Figure 2I**). Our functional pre-motor analyses therefore identify a group of five interneurons as the upstream circuit elements wired to the MN-LT1/2 system with significant impact on SR behaviour.

### Axial characterisation of the interneuronal network linked to the pre-LT1/2s

To continue our axial-structural analysis of the SR circuit, we reconstructed the upstream synaptic partners of all functionally relevant pre-motor elements described above (**Figure 3A**) along the antero-posterior axis of the larval VNC using CATMAID. This work identified the A27k, A19f and the Down-and-Back (DnB, A09l) interneurons as the upstream elements connected to the pre-LT motor neuronal components. Interestingly, previous work had described the A27k neuron as a component of the intersegmental feedback circuit underlying larval locomotion (Kohsaka et al., 2019) and the DnB neuron as a noxious heat responsive neuron involved in nociceptive behaviour (Burgos et al., 2018) illustrating the notion that neurons are often recruited to circuitries underlying distinct behaviours. The A19f has not been previously described and presents ipsilateral links to the pre-motor neuron A26f (**Figure 3A**) and, remarkably, receives input from A01c, another pre-motor element strongly suggesting that A19f might represent a pre-motor integrator (**Figure 3I**).

**Figure 3.**
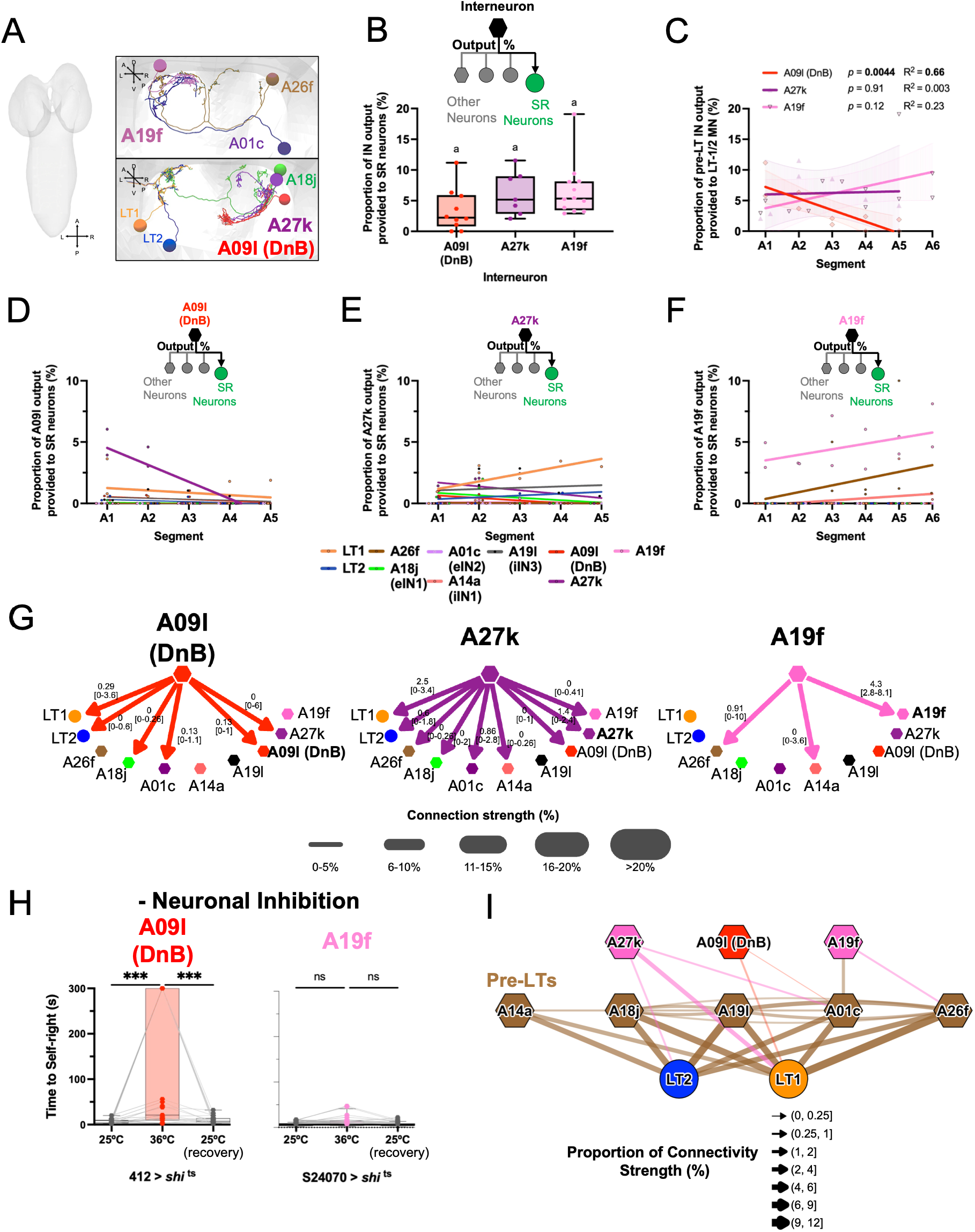
Interneurons upstream of the premotor and motor LT system along the A-P axis. **(A)** Abdominal neuronal reconstructions of interneurons upstream of the premotor and motor LT system: A09l (DnB, red), A27k (purple) and A19f (carnation). premotor interneurons (IN) of LT1/2: A26f (brown), A18j (green), A01c (purple), A14a (salmon) and A19l (black). Right panels, transverse views of the INs and the IN-MN LT system in segment A2. **(B-F)** Percentage of interneuron downstream connectivity provided to the SR neurons (upstream interneurons, pre-LTs and LT-MNs) in abdominal segments A1 to A6. (B) Boxplots with individual neuronal data points of 10 A09l (from A1 to A5), 7 A27k (A1, A2, A3 left, A4 right, A5 right), 12 A19f (from A1 to A6). Boxplots with the same letters are not significantly different (Kruskal-Wallis test with Dunn’s multiple comparisons test, n.s. *p*>0.05). (C) Bulk and (D-F) neuronal-specific percentage of IN downstream connectivity provided to SR along the abdominal segments A1 to A6 [slope n.s. *p*>0.05, \*\**p*<0.01, and goodness of fit (R^2^), simple linear regression]. (**G**) Fan diagrams displaying the downstream connectivity of each IN to the SR neurons. The thickness of the arrows represents the median downstream percentage connectivity with respective values, and minimum and maximum values in squared brackets. (**H**) SR assay and thermogenetic inhibition of the neural activity in the INs A09l (412>*shi* ^ts^; n=23 larvae) and A19f (SS24070>*shi* ^ts^; n=20 larvae). Boxplots with minimum and maximum values, and individual data points of larvae tested initially at the permissive temperature of 25°C, at the restrictive temperature of 36°C, and the recovery permissive temperature of 25°C (n.s *p*>0.05, \*\**p*<0.001, Wilcoxon matched-pairs test).

Connectomic analyses of A09l (DnB), A27k and A19f show that these interneurons display a substantial level of output connectivity to the SR neural system, with a median 3.4%, 6.2% and 6.7% of their respective total synaptic outputs linked to the SR neurons and are not significantly different among themselves (**Figure 3B**) . The axial map of connections across the abdominal segments shows a significantly marked reduction of A09l (DnB) downstream connectivity from anterior to posterior (**Figure 3C**) that is due to connections to A27k (**Figure 3D, G**). A19f has a non-significant trend of increasing SR connectivity towards posterior (**Figure 3B**) that is in part caused by connectivity to A19f INs in other segments and to A26f (**Figure 3F, G**). In contrast, A27k has a uniform pattern of axial output connectivity spread among SR neurons (**Figure 3C, E, G**).

Shi-mediated functional inhibition was conducted for A19f and A09l (DnB) and the results of these experiments show a clear role for A09l (DnB) inhibition in SR behaviour (**Figure 3H**, left) and a moderate non-significant effect on SR triggered by A19f inhibition (**Figure 3H**, right).

### The sensory elements of the SR circuit overlap with those of the rolling circuit

The discovery that the DnB (A09l) neuron – previously shown to be part of the rolling circuit (Burgos et al., 2018) – was directly linked to the pre-motor elements of the SR circuit immediately suggested the possibility that sensory inputs to DnB could play a role in SR behaviour (**Figure 4A**). In this respect, the multidendritic class IV sensory neurons (here on termed md Class IV/mdIV) (Grueber et al., 2002) emerged as an obvious candidate given their enrolment in rolling and nociceptive behaviours (Hwang et al., 2007; Ohyama et al., 2015). Analysis of md Class IV neurons outputs shows that, remarkably, more than 25% of their median synaptic output project into SR circuit elements (**Figure 4C**), with the majority targeting DnB neurons (**Figure 4E**). Axial mapping of md Class IV outputs shows a significant decline in the proportion of synaptic outputs from anterior to posterior abdominal segments (**Figure 4D**). Furthermore, *shi*-mediated inhibition of md Class IV neurons results in a significant delay in SR demonstrating a functional role of these sensory neurons in SR behaviour (**Figure H**).

**Figure 4.**
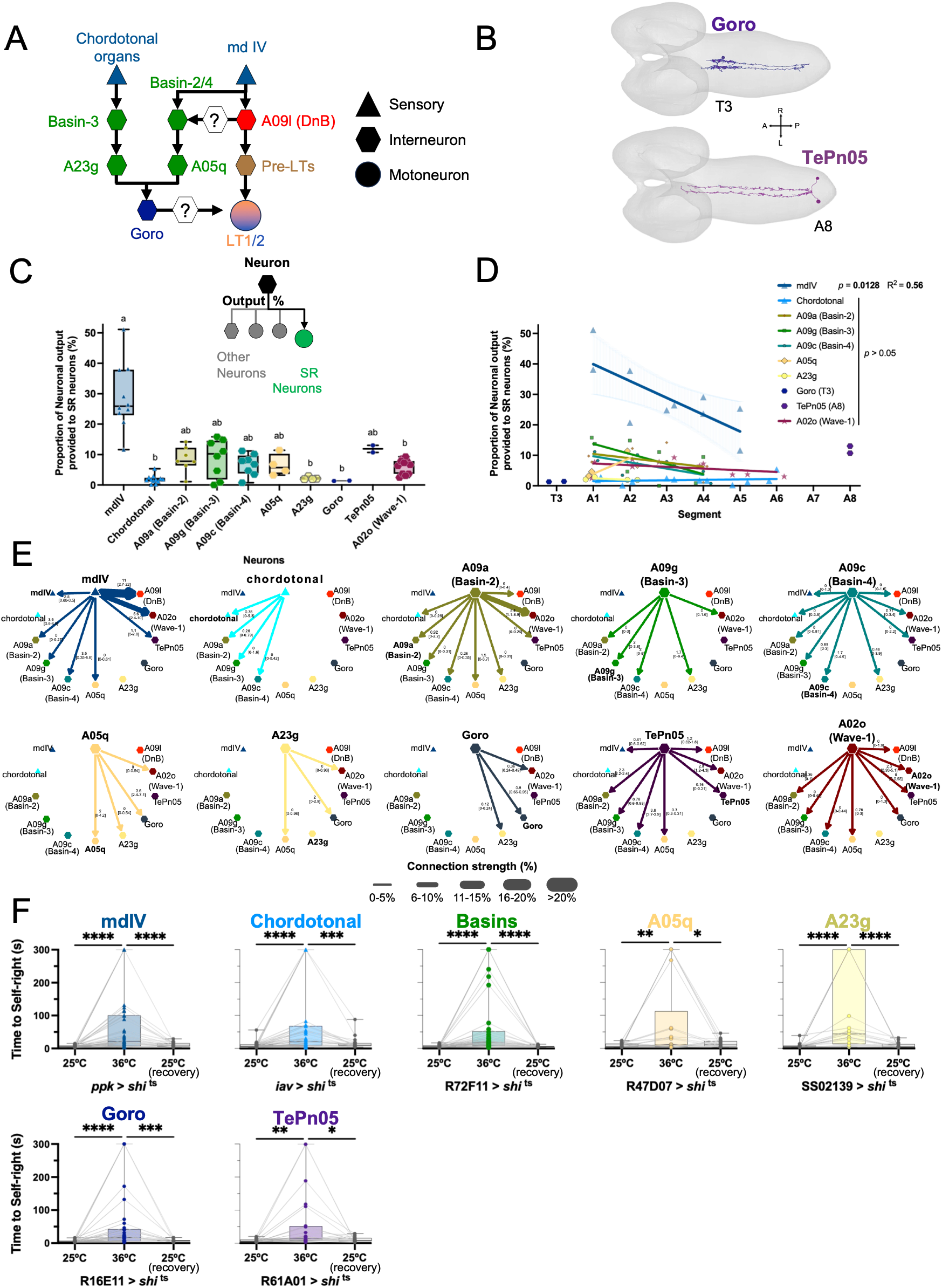
The SR circuit is connected to the rolling circuit. (A) Diagram depicting the rolling circuit, its connections to the SR circuit and missing connection links. The rolling circuit consists of the sensory neurons multidendritic class IV (mdIV) and chordotonal organs, 1° order IN basins and 2° order IN A23g and A05q, and the command-like neuron Goro. (B) First instar larval central nervous system with the neuronal reconstructions of Goro in the third thoracic segment (T3) and TePn05 in A8. **(C-E)** Percentage of downstream connectivity of rolling neurons provided to the SR neurons (rolling, DnB, TePn05 and A02o) in abdominal segments A1 to A6. (C) Boxplots with individual neuronal data points of 10 mdIV (A1 to A5), 12 chordotonal (A1-A6), 7 A09a (Basin-2, A1-A4, except A3 left), 8 of A09g (Basin-3) and A09c (Basin-4)(A1-A4), 4 of A05q and A23g (A1-A2), 2 of Goro (T3) and TePn05 (A8), and 11 of A02o (Wave-1, A1 to A5 and A6 left). Boxplots with different letters indicate significant differences (Kruskal-Wallis test with Dunn’s multiple comparisons test, n.s. *p*>0.05, \**p*<0.05, \*\**p*<0.01, \*\*\**p*<0.001). (D) Roling connectivity along the anterior-posterior axis [slope n.s. *p*>0.05, \**p*<0.05, and goodness of fit (R^2^), simple linear regression]. (**E**) Fan diagrams displaying the downstream connectivity of each rolling neuron. The thickness of the arrows represents the median downstream percentage connectivity with respective values, and minimum and maximum values in squared brackets. (**H**) SR assay and thermogenetic inhibition of the neural activity in the sensory neurons mdIV (*ppk*>*shi* ^ts^; n=40 larvae) and chordotonal (*iav*>*shi* ^ts^; n=39 larvae), Basins2,3,4 (R72F11>*shi* ^ts^; n=38 larvae), A05q (R47D07>*shi* ^ts^; n=22 larvae), A23g (SS02139>*shi* ^ts^; n=21 larvae), Goro (R16E11>*shi* ^ts^; n=34 larvae), and TePn05 (R61A01>*shi* ^ts^; n=24 larvae). Boxplots with minimum and maximum values, and individual data points of larvae tested initially at the permissive temperature of 25°C, at the restrictive temperature of 36°C, and the recovery permissive temperature of 25°C (n.s *p*>0.05, \**p*<0.05, \*\**p*<0.001, \*\*\*\**p*<0.0001, Wilcoxon matched-pairs test).

The fact that the md Class IV neurons turned out to be part of the SR circuit led us to consider the possibility that other rolling-related neurons might also be involved in SR. To test this notion, we conducted a series of functional experiments systematically inhibiting components of the rolling circuit previously described, including: sensory chordotonal organs, and first-order (Basins 2-4), second-order (A05q and A23g), and third-order interneurons (Goro) and the integrator neuron TePn05 (**Figure 4A, F)** (Burgos et al., 2018; Ohyama et al., 2015). Inhibition of each one of these neuronal elements led to a significant impact on SR put into evidence by an increase in SR time (**Figure 4F**). These findings indicate that many neuronal components are in fact part of both the SR and rolling circuits and are in line with the fact that these two behaviours, SR and rolling, involve related motor sequences that driven by distinct biological roles – i.e. postural control and escape response, respectively – concern the rotation of the larval body around its main body axis.

Connectomic analyses of output proportions suggest a high level of connectivity between Basins 2-4, A05q, TePn05 and A02o (Wave-1) neurons (median approx. 5-10%) and SR circuit neurons (**Figure 4C, E).** In contrast, chordotonal organs, A23g and Goro show very low direct connectivity to the SR circuit (median connectivity circa 1-2%; **Figure 4C, E**), suggesting that these neurons might participate in other behavioural programmes or be indirectly linked to SR components via yet unknown elements. Axial analysis of synaptic outputs for the rolling neuronal elements shows no obvious trend along the AP axis (**Figure 4D**), in contrast to the axial pattern observed in the md class IV sensory neurons.

### Integrative analysis of SR circuit properties along the antero-posterior axis

Our connectomic and functional behavioural experiments allow us to propose a first version of the SR circuit in the fly larva, which constitutes the first neural substrate for postural control mapped in this animal. Hierarchical clustering and network connectivity analyses (**Figure 5A**, B) provides the connectivity matrix for the SR circuit representing median synaptic strength amongst the individual components of the circuit. The matrix depicts the links of the sensory, interneuron and motor neuron hubs and exposes a nested topology, indicative that information is likely to flow within each hub before being transmitted upstream or downstream in the circuit. The sensory neurons md class IV show the overall highest output of any source group (∼27.4% total median output), but rather than concentrating onto one partner, its output is distributed across several Basin-associated targets and strongly to A09l (DnB). Furthermore, there are two largely parallel sub-networks: the Basin-associated group and the preLT-LT group that are connected mostly among themselves. The interneurons TePn05, A09l (DnB) and Goro act as bridging point between the two sub-networks. What’s more, the A02o (Wave-1) interneuron receives the largest total combined input of any interneuron target (∼17%), funnelled from mdIV, Basins, TePn05, Goro and DnB, making it a clear convergence SR node. These intermediate positions of A09l (DnB), A02o (Wave-1) and TePn05 that receive inputs and send output to both the sensory/Basin cluster and the LT1/LT2 base, are suggesting of an integration SR role rather than a purely feed-forward or command/-like. Interestingly, this A02o (Wave-1) interneuron has been previously described as a command-like neuron for escape behaviour by backward or forward crawling locomotion depending on whether the larvae are touched respectively on the head or on the tail (Takagi et al., 2017). This is in line with the SR behavioural sequence whereby when the larva is rectifying its postural orientation through asymmetrical rotation contracts the segments sequentially along the anteroposterior axis (Loveless et al., 2021; Picao-Osorio et al., 2015).

**Figure 5.**
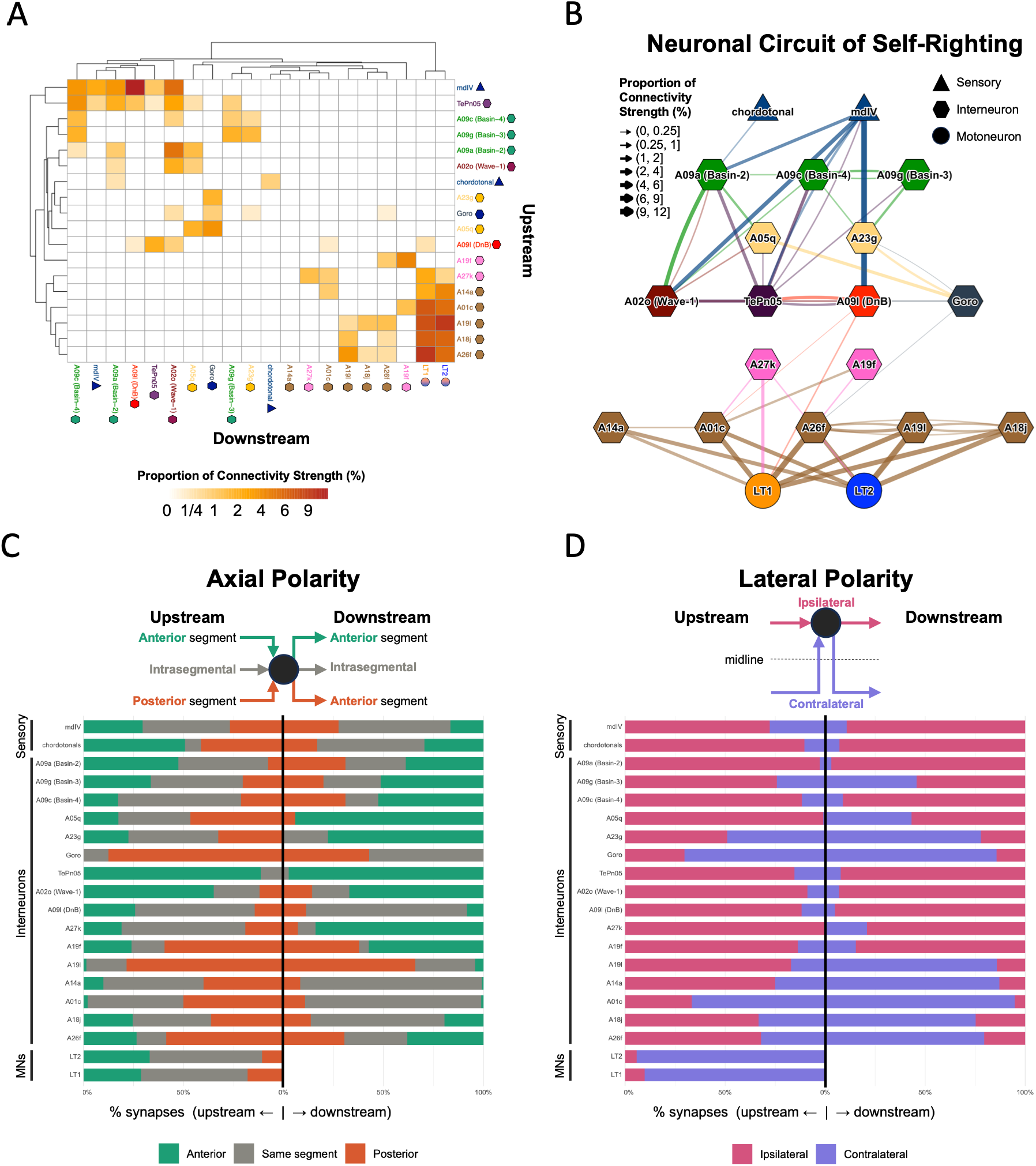
The neural wiring diagram underlying larval SR. (**A, B**) Hierarchical clustering with heatmap (A), and connectivity network (B) of the overall median proportion of downstream connections of the SR neuronal circuit. Connectivity strength is represented by darker colours in in the heatmap (A) and by edge thickness in the network (B). The SR neurons are depicted in the following way: chordotonal and mdIV sensory neurons in dark blue triangles; the first-order basin INs in green hexagons; the second-order INs in yellow hexagons (A05q and A23g); the third-order INs and integrators in midnight (Goro), maroon (A02o), aubergine (TePn05) and red (A09l/DnB); the forth-order INs in pink (A27k and A19f); the pre-LT INs in brown (A14a, A01c, A26f, A19l and A18j); and the MNs in orange (LT-1) and blue (LT-2). (C) Overall quantification of the axial connectivity polarity in the SR circuit. For a given SR neuron is shown the respective median percentage of upstream (left side) and downstream (right side) connections that come/go from/to an anterior segment (green), the same segment (grey), and a posterior segment (terracotta). (D) Overall quantification of the lateral connectivity polarity in the SR circuit. For a given SR neuron is shown the respective median percentage of upstream (left side) and downstream (right side) connections that come/go from/to an ipsilateral connection (same side, violet) and a contralateral (opposite side, medium slate blue).

Next, we asked if there is an axial and lateral connectivity polarity in the SR circuit (**Figure 5C, D**). We quantified axial polarity of each SR neuron by measuring the overall median percentage of upstream connectivity that comes from a SR neuron localised in an anterior, the same or posterior segment (**Figure 5C, left panel**). Likewise, we quantified downstream axial connectivity by measuring the median percentage of output to SR neurons localised in an anterior, the same or posterior segment (**Figure 5C, right panel**). Axial polarity was highly heterogeneous across the 21 neuron groups. It ranges from highly anterior oriented upstream and downstream connectivity of TePn05 and A02o, anterior downstream polarity in A05q, A23g and A27k, to highly posterior polarised in upstream (Goro, A19f, A19l, A26f) and downstream (A19l) connectivity. Furthermore, there are several neuronal groups dominated by instrasegmental connectivity in upstream (A27k, A09l, LT) and in downstream (A09l, Goro, and in the pre-LTs A14a, A01c, A18j). However, these axial patterns should be carefully considered, since we did not include thoracic (except for Goro) in our circuit analysis, and some neurons are only present (or were only reconstructed) in certain abdominal segments (e.g. TePn05 is only found in A8; A05q and A23g in A1-A2; A19l in A1-A3 – see Figures 2-4). Despite this, it is interestingly to observe that the integrator A02o (Wave-1) receives and send most of its SR connection from and to anterior segments, A09l (DnB) from and to the same segment, the pre-LTs A14a, A01c, A18j target SR neurons in the same segment, while the pre-LT A19l gets input and send output to posterior segments.

We quantified lateral polarity of each SR neuron by measuring the overall median percentage of connections from the same side (ipsilateral) and from the opposite side (contralateral) in reference to the midline in upstream (**Figure 5D, left panel**) and downstream (**Figure 5D, left right**) neurons. Lateral polarity is highly skewed with most sensory and interneural components of the circuit receiving and sending information ipsilaterally. In contrast, the SR/rolling neurons A23g and Goro have a contralateral bias in their upstream and downstream connectivity, and the pre-LT A01c and MN LT1 and LT2 receive most of their input from contralateral connection. In line with the contralateral innervation of the MN-LT1/2s, the pre-LTs A19l, A14a, A01c, A18j, A26f have most of their downstream connections to neurons on the opposite side of the mideline. This observations suggest a highly integrated system for the relay of information between left and right aspects of the circuit. This connectivity pattern is well aligned with the features of SR behaviour which implies a coherent integration of sensory motor patterns to successfully initiate and complete the postural control sequence.

## CONCLUSION

This work provides the first comprehensive wiring diagram of the neural circuit underlying self-righting in the *Drosophila* larva, extending from the sensory periphery through several layers of interneurons to the LT1/2 motor neurons (Picao-Osorio et al., 2015; Zwart et al., 2016) across the full antero-posterior extent of the abdominal segments rather than in a single segment in isolation. By combining connectomic reconstruction (Ohyama et al., 2015; Schneider-Mizell et al., 2016) with systematic functional neuronal inhibition (Brand and Perrimon, 1993; Kitamoto, 2001), we show that the great majority of the identified components — from the pre-motor interneurons A18j, A01c, A14a and A26f through to the DnB, mdIV and rolling-circuit elements — make a genuine behavioural contribution to SR, rather than being connectivity artefacts of the EM volume. The overlap we uncover between the SR, rolling and crawling circuits, most strikingly at the level of the mdIV sensory neurons and the DnB, A02o (Wave-1) and TePn05 interneurons (Burgos et al., 2018; Ohyama et al., 2015; Takagi et al., 2017), indicates that the larval nervous system reuses a common pool of sensory-motor infrastructure to support distinct, and perhaps at times competing, behavioural outputs, with the specific pattern of recruitment and axial weighting presumably determining which motor sequence is ultimately executed.

Beyond the identity of the individual neurons, the network, axial and lateral connectivity analyses point to a set of organisational principles that may generalise beyond this particular circuit: a nested, hub-like topology in which information is processed locally before being relayed onward; a small number of neurons — A02o (Wave-1), A09l (DnB) and TePn05 in particular — acting as integrators bridging sensory and motor sub-networks; and a marked antero-posterior gradient in both synapse number and proportion connectivity in MNs-LT-1/2, the pre-LT A01c, the integrator interneuron A09l/DnB and the sensory md class IV neurons that could plausibly account for the segmental differences in SR performance reported previously (Loveless et al., 2021; Picao-Osorio et al., 2017, 2015) (Picao-Osorio et al. 2015; Loveless et al. 2021). We are conscious that several elements of the circuit were only reconstructed in a subset of segments, and that our analysis is necessarily restricted to a single, first-instar EM volume (Ohyama et al., 2015; Schneider-Mizell et al., 2016) so the axial trends we describe should be treated as a first approximation rather than a definitive map. Nonetheless, this circuit offers a concrete, testable scaffold for future work — whether to dissect the precise computations performed at each integrator node, to ask how the Hox-regulated microRNAs identified in earlier work act on this specific wiring (Mallo and Alonso, 2013; Picao-Osorio et al., 2017, 2015), or, more speculatively, to inform the design of axially distributed control architectures in soft-bodied robotics (Loveless et al., 2021).

## ACKNOWLEDGEMENTS

We thank past and current members of the Alonso lab for helpful discussions and comments, especially Edward O’Garro-Priddie for all his contributions to this study. We thank Marta Zlatic, James W. Truman, Matthias Landgraf and the Bloomington Stock Centre for fly stocks. The authors thank HHMI Janelia Research Campus for funding and support, and the HHMI Janelia Visiting Scientist programme to CRA for providing critical training in EM connectomics. This research was funded by a UK Wellcome Trust Investigator Award (098410/Z/12/Z), UK Medical Research Council Project Grant (MR/S011609/1) and a UK Biotechnology and Biological Sciences Research Council Project Grant (BB/Y006860/1) awarded do C.R.A.; Core funding from HHMI JRC and MRC LMB, and a UK Wellcome Trust grant (205038/Z/16/Z) to A.C.. J.P.-O. is funded by a EU Marie Skłodowska (101110355-EvoBias), a Portuguese Exploratory Grant by Fundação para a Ciência e Tecnologia (FCT) (OrgBias/2023.11630.PEX) and the FCT strategic project UIDB/00329/2020 granted to CE3C. We extend our gratitude to everyone whose work contributed to reconstructing the larval abdominal circuit, whether directly for this paper or through foundational past publications: E. O’Garro-Priddie, M. Zwart, T. Ohyama, S. Takagi, A. Burgos, A. Fushiki, C. Schneider-Mizell.

